# *Toxoplasma gondii* infection triggers chronic cachexia and sustained commensal dysbiosis in mice

**DOI:** 10.1101/247866

**Authors:** Jessica A. Hatter, Yue Moi Kouche, Stephanie J. Melchor, Katherine Ng, Donna M. Bouley, John C. Boothroyd, Sarah E. Ewald

## Abstract

*Toxoplasma gondii* is a protozoan parasite with a predation-mediated transmission cycle between rodents and felines. Intermediate hosts acquire *Toxoplasma* by eating parasite cysts which invade the small intestine, disseminate systemically and finally establish host life-long chronic infection in brain and muscles. Here we show that *Toxoplasma* infection can trigger a severe form of sustained cachexia: a disease of progressive weight loss that is a causal predictor of mortality in cancer, chronic disease and many infections. *Toxoplasma* cachexia is characterized by acute anorexia, systemic inflammation and loss of 20% body mass. Although mice recover from symptoms of peak sickness they fail to regain muscle mass or visceral adipose depots. We asked whether the damage to the intestinal microenvironment observed at acute time points was sustained in chronic infection and could thereby play a role the sustaining cachexia. We found that parasites replicate in the same region of the distal jejunum/proximal ileum throughout acute infection, inducing the development of secondary lymphoid structures and severe, regional inflammation. Small intestine pathology was resolved by 5 weeks post-infection. However, changes in the commensal populations, notably an outgrowth of *Clostridia spp*., were sustained in chronic infection. Importantly, uninfected animals co-housed with infected mice display similar changes in commensal microflora but never display symptoms of cachexia, indicating that altered commensals are not sufficient to explain the cachexia phenotype alone. These studies indicate that *Toxoplasma* infection is a novel and robust model to study the immune-metabolic interactions that contribute chronic cachexia development, pathology and potential reversal.

## Introduction

Chronic diseases account for over 85% of deaths in the first world and 70% of deaths globally(1). The co-occurrence of cachexia, or the progressive loss of lean body mass, is one of the best predictors of mortality across chronic disease. Cachexia is distinct from starvation or malabsorption and can be accompanied by anorexia, elevated inflammatory cytokines (IL-1, IL-6 and TNF), loss of fat and insulin resistance(2). In human disease, therapeutic interventions including nutritional supplementation, appetite stimulants, steroid treatment and TNF inhibitors have not proven widely successful to block or reverse cachexia(3). Current animal models of cachexia are limited in that they are transient (endotoxin injection) or have a short window of study between weight loss onset and death (cardiac, gastric or renal obstruction surgeries and tumor)(4). There is a great need to develop experimental tools to study the biology of chronic cachexia and identify targets for disease intervention.

*Toxoplasma gondii* is an obligate intracellular protozoan parasite that cycles between a broad range of mammalian intermediate hosts and definitive feline hosts. Intermediate hosts are infected for life and support haploid division/asexual expansion of parasite strains. Intermediate hosts are infected when they ingest oocysts shed cat in feces or tissue cysts, termed bradyzoites, from muscle or brain of other intermediate hosts. Over the first three days post-ingestion, *Toxoplasma* migrates down the small intestine, converting to the rapidly dividing tachyzoites stage, infecting intestinal epithelial cells and immune cells(5–7). Acute infection is marked by severe, focal disruption of the villi, expansion of secondary lymphoid structures and the appearance of “casts” formed from matrix and dead cells that form a physical barrier over damaged regions of the ileum(7). Several groups have reported a decrease in microbial diversity in the gut, an outgrowth of Gram negative bacterial species, as well as commensal microbe translocation to the liver(7–9). However, whether these alterations to commensal homeostasis are maintained during chronic infection has not been asked.

*Toxoplasma* benefits from local intestinal inflammation by infecting infiltrating monocytes and dendritic cells and using them to traffic throughout the host(10). Over the course of three to four weeks, a Th1-mediated adaptive immune response clears systemic parasitemia, except in immune-privileged tissues (mainly the brain and skeletal muscle) which support stage conversion to bradyzoite tissue cysts. Bradyzoites are characterized by altered transcriptional profiles, a shift to glycolytic metabolism, slow growth and formation of a polysaccharide-rich wall that protects the parasites as they transit through the stomach of the subsequent host(11). Thus, parasite transmission requires a robust host immune response; ensuring that the host survives acute infection and enabling the parasite to access the tissues amenable for chronic infection. Yet, once the parasite has converted to the bradyzoite, transmission requires predation of the chronically infected host. Cats acquire *Toxoplasma* by eating intermediate hosts and play an important role in the parasite life cycle by: 1) facilitating sexual recombination of the parasite thereby increasing genetic diversity; and 2) mediating range expansion of the parasite by shedding millions of highly stable and highly infectious oocysts(12,13). The selective advantage conferred by infecting cats and the predator-prey relationship between cats and rodents suggest that mice and rats are critical intermediate hosts for *Toxoplasma*. The importance of this relationship is evident in the sophisticated mechanisms the parasite has evolved to intersect host signaling pathways(14), promoting intracellular survival; as well as the observation that *Toxoplasma* infected rodents lose their aversion to cat urine, a putative means to facilitate transmission(15,16).

Here we show that in the first 10 days post-*Toxoplasma* infection adult mice lose 20% of their body mass, associated with elevated circulating cytokines, anorexia and moribund behavior. The majority of *Toxoplasma* infected animals do not succumb to infection yet the reduction of muscle mass and visceral white adipose depots is sustained. We show that *Toxoplasma* infects and replicates in distinct puncta in the distal jejunum and proximal ileum throughout the acute phase of infection. Peak inflammation correlates directly with parasite load but is resolved by 5 weeks post-infection. Using 16S sequencing, we identify an outgrowth of *Clostridia spp.* that is sustained during the chronic stages of disease. Importantly, co-housed uninfected animals exhibit a similar shift in commensal populations without exhibiting any signs of illness or weight loss, consistent with the conclusion that commensal alterations alone is not sufficient to explain the sustained cachexia disease. We propose that promoting muscle and fat wasting may be a means of enhancing the opportunity for rodent predation and transmission of this parasite to definitive feline hosts.

## Materials and Methods

### Animals

CBA/J, BALB/cJ, C57BL/6J and 1291/SvImJ mice were purchased from Jackson Laboratories. Animals were housed in BSLII level conditions. All animal protocols were approved by Stanford University’s Administrative Panel on Laboratory Animal Care (Animal Welfare Assurance # A3213-01, protocol # 9478) or The University of Virginia Institutional Animal Care and Use Committee (protocol # 4107-12-15) All animals were housed and treated in accordance with AAALAC and IACUC guidelines at the Stanford School of Medicine or the University of Virginia Veterinary Service Center.

### Parasites, cells and cell lines

The parasite strain used for these studies was Me49 that stably expresses green fluorescent protein and luciferase, and has been previously described (17). Parasites were passaged intracellularly in human foreskin fibroblasts (ATCC) and passaged by 25G syringe lysis in complete DMEM (cDMEM, Gibco) plus 10% FBS (HiClone), 100ug Penicillin-Streptomycin (Gibco) and 1mM Sodium Pyruvate (Gibco).

### Infections

To generate cysts, 6-8 week-old female CBA/J mice were infected with 1000 Me49 tachyzoites stably expressing green fluorescent protein and luciferase (Me49-gfp-luc) by intraperitoneal injection. 4-8 weeks following infection, mice were euthanized with CO2 and brains were harvested, homogenized through a 50 μm filter, washed 3 times in PBS, stained with dolichos biflorus agglutinin-rhodamine (Vector labs) and the number of cysts were determined by counting rhodamine GFP double-positive cysts at 20x magnification. Prior to infection, 8-10 week male mice were cross-housed on dirty bedding for two weeks to normalize commensal microbiota. Mice were fasted overnight and fed between 100 and 250 Me49-GFP-luc cysts on ¼ piece of mouse chow. Weights and health were monitored daily. To measure food intake, mice were house on chip bedding and food was weighed daily and normalized to totally mouse body weight in the cage.

### Tissue harvesting and cytokine measures

At the experimental end points, mice were euthanized with CO2. Blood was isolated by cardiac stick, abdominal sub cutaneous white adipose depots, epidedimal visceral white adipose depots, neck brown adipose depots, quadriceps, tibialis anterior, EDL and gastrocnemius muscles were isolate and placed in pre-weighed 2mL tubes for weighing. For small intestine, a Peyer’s Patch containing regions of the distal jejunum or ileum were identified by eye. A 2cm section surrounding the Peyer’s patch (or patches) was excised. Sections immediately adjacent to but excluding a Peyer’s patch were harvested as well. Sera cytokines levels were measured by Luminex cytokine array at the Stanford Human Immune Monitoring Core or at the University of Virginia Flow Cytometery Core.

### Bioluminescence imaging and quantification

For bioluminescence imaging (BLI), mice were injected in the intraperitoneal cavity with with 200 μL of a 15 mg/mL stock solution of luciferin (Xenogen), anesthetized with isoflurane and imaged for 4 minutes on an IVIS system. To image organs, mice were injected 5 minutes prior to euthanasia, organs were harvested and imaged for 4 minutes. Images were analyzed with LivingImage software and ImageJ.

### Histology and microscopy

At the experimental end points, mice were euthanized with CO2, tissues were harvested and subjected to IVIS imaging and fixation in formalin. Samples were submitted to the Stanford Department of Comparative Medicine Histology Core for paraffin embedding and sectioning. For regions of the jejunum were selected based on the presence of a Peyer’s patch. Adjacent sections were taken and every other section was stained with H&E. A semi quantitative scoring system of 1 to 5, (1 = no significant lesion, 2 = mild, 3 = moderate, 4 = marked, 5 = severe) was used to evaluate the severity of any lesions. Parameters included inflammatory cellular infiltrate, loss of Peyer’s patch organization, villi destruction and villi shortening. For detailed scoring, each tissue section was divided into fields of view at 40x and an inflammation score was assigned to each field of view.

Unstained sections were used to quantify *Toxoplasma* load. To deparaffinize, sections were passed twice through xylene, then through 100% ethanol, 80% ethanol and 50% ethanol and distilled water for 3 minutes each. Antigen retrieval was performed by incubating sections in sodium citrate buffer brought to a boil in the microwave and incubated for 15 minutes in a vegetable steamer (10mM Citric Acid, 0.05% Tween-20, pH6.0). Slides were cooled, washed once in PBS and outlined with a Pap pen to perform staining. Samples were blocked for 30 minutes in 5% goat sera in PBS. *Toxoplasma* was labeled with mouse anti-*Toxoplasma*-FITC (Thermo Scientific Clone J26A) at a concentration of 1 μg/μL in 5% goat sera overnight. Samples were washed 3x in PBS, mounted in Vectashield with DAPI (Vector Laboratories) and imaged on an Olympus BX60 upright fluorescence microscope with a 4x, 10x, 40x or 100x objective. To quantify parasite load, each section was subdivided identically to the adjacent H&E section. The threshold of parasite signal at 488nm was determined by comparison to uninfected samples, each image was converted to binary and the dark pixels were counted using ImageJ.

### 16S ribosomal sequencing and diversity analysis

Fresh fecal pellets were collected from mice at the time points indicated and flash frozen. DNA was isolated using the MoBio PowerSoil Kit and bar coded primers were used to amplify the V4 region of the 16S rRNA gene. MoBio UltraCLean-htp 96 Well PCR Clean-Up Kit was used to purify PCR products which were then quantified using IQuant-iT ds DAN Assay Kit. 184 samples were pooled at equimolar ratios. 16S ribosomal sequencing was performed by the Mayo Clinic using a single lane of the Illumina HiSeq. Community composition and beta diversity were determined using QIIME and beta diversity was visualized using EMPeror(17,18). T-tests were performed using GraphPad Prism and corrected for multiple hypothesis using the FDR approach.

## Results and discussion

10-12 week male C57BL6 mice acquired from Jackson Labs were cross-housed for two weeks then infected per orally with 100-250 *Toxoplasma* cysts of the Me49 background engineered to express GFP and luciferase (Me49-GFP-luc). Body weight was monitored for the duration of infection. Mice lost a significant amount of weight during the acute phase of infection, 7 to 14 days post infection (dpi), but weight loss stabilized by 30 dpi, the onset of chronic infection (Fig 1A). Although mice increased in weight over the chronic phase of infection (day 30-90) they remained 20% less massive than uninfected controls. Animals infected with 100 or 250 cysts had similar weight loss (Fig 1B) and survival through 40 dpi (Fig 1C). As expected, parasites were visible by bioluminescence assay when imaged ventrally at day 7 dpi (Fig 1D); however, by day 40 dpi, parasite signal was not detectable (Fig 1E). Of note, this phenotype was not restricted to C57BL/6 mice. CBA/J mice, also lost approximately 20% of their body mass; however, BALB/C mice which have a protective H-2L^d^ haplotype do not exhibit weight loss in response to *Toxoplasma* infection (Fig F)(19,20). Although mice underwent a phase of anorexia during acute infection, they regained their appetites and consumed normal pre-infection food amounts by 15dpi indicating that sustained weight loss was not simply due to anorexia (Fig 1G). These results are consistent with a 1997 report from Arsenijevic et al. that showed that mice infected with *Toxoplasma* can be described in different response groups: death in acute infection, failure to regain body mass or partial recovery of body mass(21). However, the physiological basis for this weightloss was not determined. To identify the tissue types effected, abdominal subcutaneous fat depots (scWAT, a rapidly mobilized energy source), epidedimal visceral white adipose depots (vWAT, a key metabolic regulatory tissue), and supraclavicular brown adipose depot (BAT, thermogenic fat) were harvested. At 10 dpi, there was already significant reduction in BAT and scWAT depots. VWAT depots were significantly reduced 5 weeks post infection (wpi) (Fig 1H). In contrast to fat depots, which were reduced early, tibialis anterior (TA), gastrocnemius (GA) and quadriceps (QUAD) muscles were significantly reduced at 5wpi (Fig 1I). This sustained muscle loss was consistent with the recent observation of T reg mediated myositis during chronic *Toxoplasma* infection which leads to impaired animal strength(22). At 7 dpi, the canonical cachexia cytokines IL-1β, TNF-α and IL-6 were significantly upregulated in the sera (Fig 1J). Some inflammatory cytokines were still detected 5wpi, although the overall magnitude of inflammation was greatly reduced. Cumulatively these data indicate that chronic infection with *Toxoplasma* meet a modern definition of cachexia put forth in 2008: the loss of 5% or more lean body mass accompanied by anorexia, fat loss, inflammation (IL-1, TNF, IL-6, acute phase proteins) and/or insulin resistance(2).

**Fig 1.**
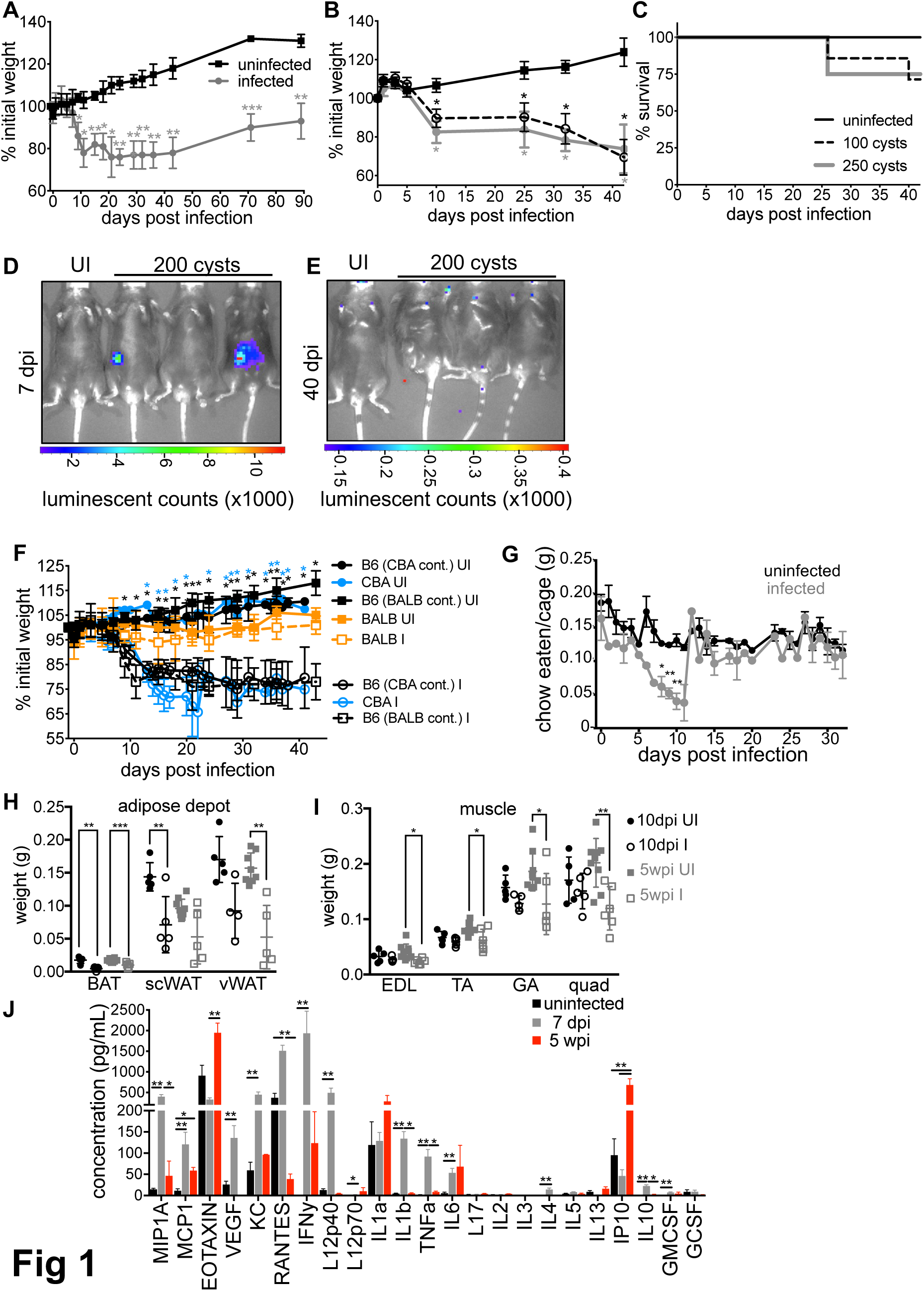
C57BL/6 mice infected with *Toxoplasma* become chronically cachexic. (A) Following per oral infection with 120 Me49-GFP-luciferase cysts (grey) or mock infection (black) mice were monitored for weight loss. Data average of 8 experiments. (B) Mice were infected as described in A with 100 cysts (dashed line), 250 cysts (grey) or mock infected (black) weight was monitored at the indicated time points. (C) Survival curves for mice represented in B. N=4-7 mice per group, data representative at least 3 experiments. (D) Mice were harvested at 7 days post infection (dpi) (E) or 40 dpi to assess parasite load by bioluminescence assay. (F) BALB/c (orange square) or C57BL 6/J mice (black square, B6 BALB cont.); CBA/J (blue circles) or C57BL 6/J (black circles, CBA cont.) were infected with 120-200 cysts or mock infected. Weight was monitored at indicated time points. N=4-8 mice per condition averaged across two independent experiments, significance is measured relative to uninfected at same time point. (G) Food was weighed every 24 hours to determine the amount eaten and normalized to the weight of animals in the cage. Data are average of 2 experiments, N=5 mice per group each experiment. Significance relative to uninfected at the same time point. H, Brown adipose tissue (BAT), sub cutaneous white adipose tissue (scWAT) or visceral white adipose tissue (vWAT) was harvest at 10 dpi (black) or 5 wpi (grey) and weighed. I, Extensor digitorum longus (EDL), tibialis anterior (TA), gastrocnemius (GA) and quadriceps (QUAD) muscles were weighed at 10 dpi (black) or 5 wpi (grey) post infection. Data is average of 2 experiments, N=5-10 mice per time point. J, Luminex cytokine array was performed on sera from uninfected mice (black), 7dpi (grey) or 5 wpi (red) mice. N=3-10 mice per group. * p≤0.01, ** p≤0.001, ***p≤0.001, SEM, student’s T-test.

*Toxoplasma* is naturally acquired by ingestion of oocysts or tissue cysts leading to severe regional inflammation in the small intestine, a breakdown in small intestinal architecture and interaction with commensal microbiota that drive a TLR-mediated innate immune response (7,9,23). We reasoned that cachexia could be the result of sustained inflammation, changes in intestinal architecture and/or the gut commensal community. To address this question, we orally infected mice with 200 Me49-GFP-luc cysts. Three mice per day were euthanized to assess parasite load in the small intestine, mesenteric lymph node and spleen by bioluminescence assay. Significant parasite signal was observed in the small intestine at 4 dpi and peaked 7-8 dpi (Fig 2A, black bars), preceded by parasite dissemination to the mesenteric lymph nodes (Fig 2A, green bars) and spleen (Fig 2A, grey bars). For as long as *Toxoplasma* was detected by BLI (Fig 2A, 4-10dpi) the first luciferase signal was consistently found at 50% the length of the small intestine and the mean of all luciferase positive regions was identified at 2/3^rd^ the length of the intestine (Fig 2B). Gregg and colleagues have shown parasite infection along the mucosa of the small intestine in the duodenum, jejunum and ileum in the first 6 days of infection(6). Further, Molloy et al. demonstrated that 9 dpi, commensals were segregated from the epithelial layer in the ileum but not the jejunum by the presence of a ‘cast’-like pseudomembrane composed of dead host cells and invasive *E. coli* suggesting that there is a distinct interplay between *Toxoplasma*, commensals and the immune system in this tissue(7). While we did not observe parasite signal in the duodenum, this may be due the fact that we imaged intestines from the serosal side rather than the luminal aspect. In addition bioluminescence assay is not sensitive enough to detect small numbers of parasites that may be present elsewhere in the small intestine(6). None the less, our data are consistent with the interpretation that the distal jejunum/proximal ileum is the major small intestinal niche for parasite replication throughout acute infection. This region of the small intestine is enriched in immune resident cells, specialized structures, like M cells that allow for sampling of the lumen and pathogen transit as well as an expansion in microbial diversity any of which could contribute to *Toxoplasma’s* predilection for residence in this niche(24).

**Fig 2.**
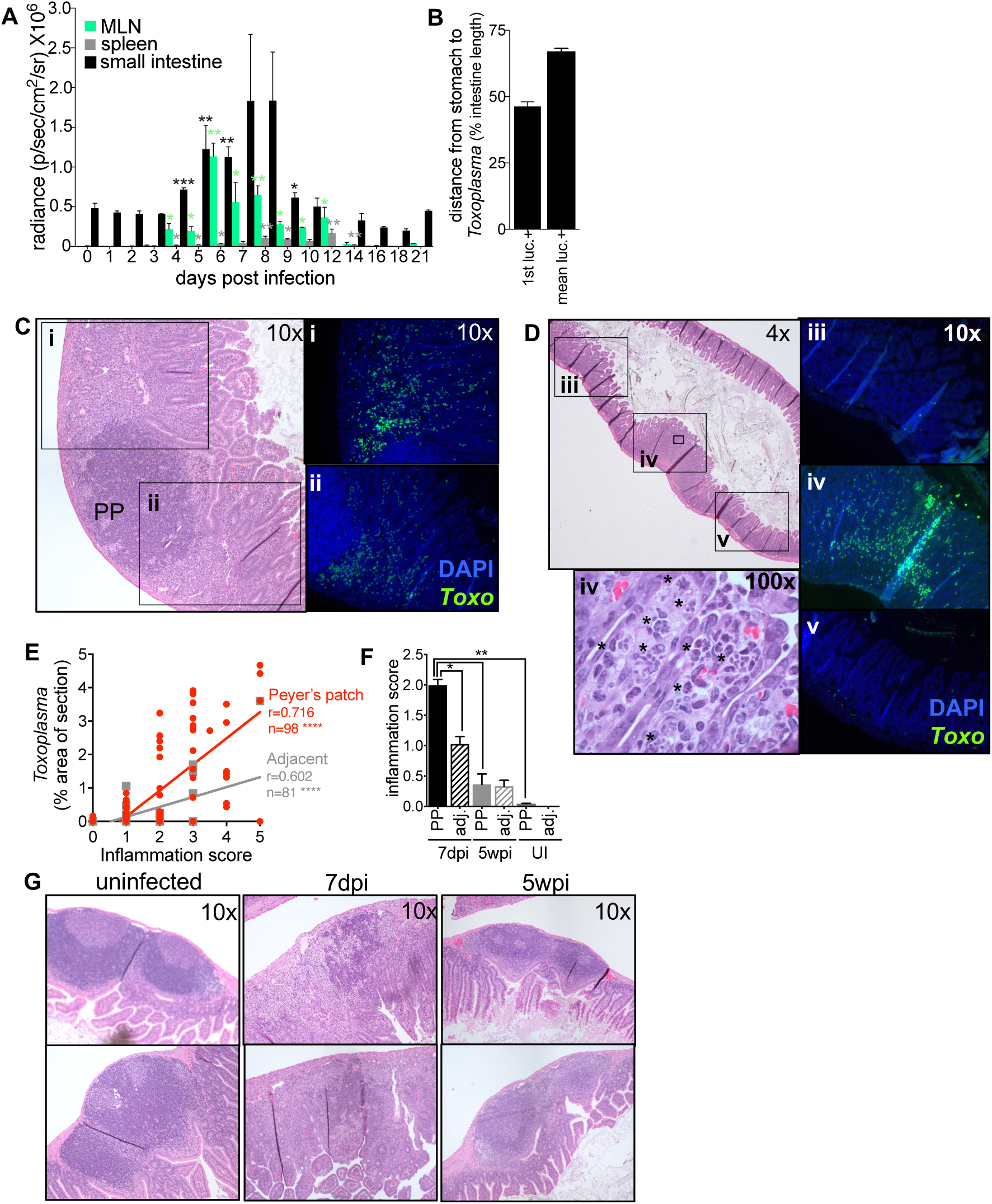
Mice recover from severe acute inflammation and parasite growth in the small intestine. (A-B) Following per oral infection with 200 Me49-GFP-luc cysts, mice were euthanized and parasite load was determined by BLI in the small intestine (black), mesenteric lymph nodes (MLN, green) and spleen (grey). N=3 animals per day, Significance is measured for each time point relative to day 3 post infection. Student’s T-test * p≤0.01, ** p≤0.001. (B) For each intestine, the position of the luciferase signal was measured as distance from the stomach to the first luciferase positive region (1^st^ luc+) or mean of all luciferase positive regions (mean luc +). Position was averaged across 4 to 10 dpi. (C-H) 2cm segments of the distal jejunum/ileum (the distal 50-90% of the small intestine) were excised for histology based on the presence of Peyer’s patches visible to the eye at 7dpi, 5wpi or from uninfected mice.(C, D) Representative images of 7dpi sections stained with H&E to assess inflammations score at 10x magnification (scale of 1=no detectable inflammation to 5=complete disruption of lymphoid structure and/or villi) or to identify parasite vacuoles at 100x magnification (iv, asterisks). An adjacent section of each intestinal segment was stained with a *Toxoplasma*-specific antibody (green) and DAPI (blue and imaged at 10x (i-iv) to assess parasite load. (E) Strength of the correlation between parasite load and inflammation score for 7dpi intestine segments containing a Peyer’s patch (red) or adjacent segments lacking a Peyer’s patch (grey), Pearson’s correlation **** p≤0.0001. Slopes were significantly different between the two groups: Peyer’s patch 0.781±0.078, adjacent 0.296±0.044 p<0.0001, linear regression of correlations. (F) Inflammation score for Peyer’s patch-containing segments (solid bars) and adjacent, Peyer’s patch negative segments (dashed bars) of the intestine in uninfected animals, 7 dpi (black) or 5wpi (grey). Student’s T-test *p≤0.05, ** p≤0.001, SEM, N=4-6 segments from 3 mice per condition. (G) Representative images of Peyer’s patch organization in uninfected 7dpi and 5wpi small intestines.

Diet as well as reactive oxygen species derived from inflammatory infiltrate can produce auto-luminescent signal. To validate that the luciferase signal was derived from *Toxoplasma*, and to monitor the degree of inflammation, we harvested segments of the small intestine 7 dpi for histological analysis. Having observed that luciferase positive regions always occurred adjacent to at least one enlarged Peyer’s patch, 2cm segments centered on a Peyer’s patch (or Peyer’s patches) were excised from the small intestine of infected and uninfected mice (Fig 2C-G). This allowed us to assess parasite load and degree of inflammation in matched regions of the intestine across time points without pre-existing knowledge about parasite location provided by BLI. These Intestine segments were fixed and sectioned. One section was stained with H&E (Fig 2C, D, G) to assess inflammation. The adjacent section was de-paraffinized and stained for *Toxoplasma* using an antibody specific to parasite lysate and for nuclei using DAPI (Fig 2i-iv). At 7 dpi, tachyzoites were observed throughout the villi and the lamina propria. Interestingly, in sections where a Peyer’s patch was cross-sectioned, parasites were observed nearby but excluded from lymphoid follicle (Fig 2Ci and ii). When H&E sections were examined at high magnification, vacuoles containing multiple tachyzoites were visible in intestinal epithelial cells, indicating that parasites were growing in this niche at 7dpi (Fig 2Div, 100x, asterisks).

We noticed the fields of view closest to the Peyer’s patch contained the most *Toxoplasama* (Fig 2D iv), whereas neighboring fields of view contained few parasites and were less morphologically disrupted (Fig 2D iii and v). To quantify this observation, 2cm segments of the intestine containing a Peyer’s patch or 2cm segments immediately adjacent to but excluding Peyer’s patches were isolated, sectioned, and stained for H&E or *Toxoplasma* as described above. Across each section, there was a significant positive correlation between inflammation score and parasite load (Fig 2E Pearson’s correlation, Peyer’s patch segments, red: r=0.716, n=98, p<0.0001; adjacent segments, grey: r=0.602, n=81, p<0.0001). Peyer’s patch negative sections had a significantly lower overall inflammation score and parasite load (Fig 2E linear regression of correlations, Peyer’s patch segments: 0.781±0.078; Adjacent segments: 0.296±0.044, p<0.0001 and Fig 2F). By 5 wpi there was no significant difference between parasite load or inflammation score in infected versus uninfected intestinal segments (Fig 2F). Also consistent with the conclusion that infection in the small infection was resolved in chronic disease, Peyer’s patch architecture, which had a highly disorganized germinal center 7 dpi, was indistinguishable from uninfected animals by H&E staining 5wpi (Fig 2G). Taken together, these results indicate that acute inflammation in the small intestine is resolved by chronic infection and is therefore unlikely to drive the sustained cachexia in these animals.

Several groups have observed that acute infection with *Toxoplasma* triggers a loss of microbial diversity, an enrichment in Gram negative bacteria associated with intestinal pathology, and, sometimes, lethal ileitis(7,9,25). However, it is not known if these changes in the commensal communities are transient or sustained in chronic infection. To address this, we collected fecal pellets from mice over the course of infection with *Toxoplasma* and analyzed microbial diversity by 16S ribosomal sequencing (Fig 3). In each cage, 1-2 uninfected animals were co-housed with infected littermate controls. Community composition analysis reflected significant expansion in *Clostridia spp*. OTUs 5 weeks post-*Toxoplasma* infection when compared to the pre-infection community (Fig 3A navy blue outset, 5 wpi 0.599±0.142 versus pre-infection 0.147±0.015 SEM, p=0.0004, q=0.007 student’s T-test). This trend was also observed in uninfected animals, although it was not statistically significant (5wpi 0.373±0.181 versus pre-infection 0.100±0.005 SEM, p=0.319, q=0.569). There was also a moderate, expansion in Verrucomicrobia at 1 week in both infected and uninfected animals that contracted by 5 weeks, although this change was not significant. The enrichment of *Clostridia spp*. in fecal pellets of infected mice was unexpected, based on previous observations that *Toxoplasma* infection can trigger an outgrowth of γ-proteobateria in the lumen of the small intestine 7-9 dpi (7,8,25,26). When interpreted in the context of previous reports, these results are consistent with a model where the outgrowth species may reflect the facultative pathogens already present in the community (dependent on mouse genetic background and facility to facility variation) that capitalize on niche availability following an inflammatory insult, rather than a specific relationship between a specific facultative species and *Toxoplasma*. Interestingly, enrichment in *Clostridia spp*. has been associated with expanded populations of regulatory T cells in the intestine, which may explain why our mice are more resistant to a high dose infection with Me49 cysts than others have reported in the past (9,21,24). A second parameter that may explain why our mice are tolerant of high dose infection is that uninfected and infected animals were co-housed for the duration of the study. Coprophagia may have buffered the severity of the commensal shift in infected mice as well as altered the commensal population in uninfected animals, as discussed below.

**Fig 3.**
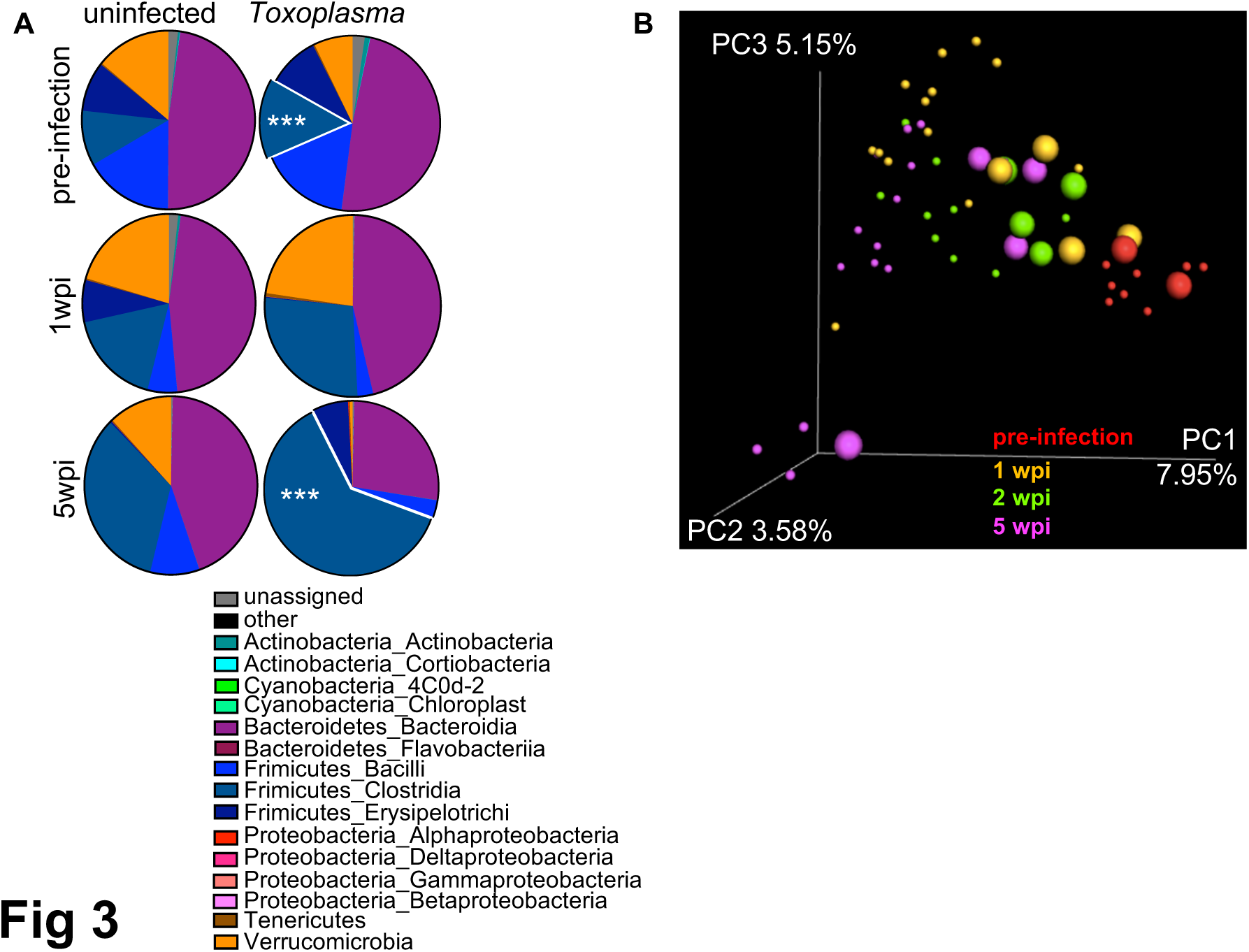
Changes in the commensal community are amplified in chronic infection. 16S profiling of fecal pellet commensal microbiota before pre-infection, 1wpi or 5wpi with 120 *Toxoplasma* cysts. (A) At chronic infection there is a significant outgrowth of *Firmicutes Clostridia* in comparison to pre-infection (navy blue, outset). *Toxoplasma* infected group: pre-infection mean 0.147±0.015 SEM, 5wpi mean 0.599±0.142 SEM, ***p≤0.001 student’s T-test. Data are mean of 3-8 mice per time point. (B) Unweighted Unifrac principle component analysis of 16S ribosomal subunit diversity in the fecal pellets of mice pre-infection (red), 1 wpi (orange), 2 wpi (green) or 5 wpi (magenta). Small circles=infected, large circles = uninfected. Data representative of two experiments.

When principal component analysis was used to assess beta diversity across fecal pellets, pre-infection animals (Fig 3B red) clustered distinctly from infected animals (Fig 3B small circles: yellow=1 wpi, green=2 wpi; magenta=5 wpi). Interestingly, the co-housed uninfected animals had a shift in microbial diversity as well and represented an intermediate cluster between pre-infection and infected animals (Fig 3B, 1 wpi yellow, large circles; 2 wpi green, large circles). The shift in similarity away from pre-infected phenotype was most pronounced by 5 wpi. (Fig 3B magenta, small circles=infected, large circles=uninfected). As co-housed, uninfected animals do not display symptoms of cachexia we conclude that the observed changes to the microbial species are not sufficient to explain the cachexia phenotype alone. However, future studies will be needed to understand if the altered commensal community synergizes with immune or metabolic defects to promote cachexia maintenance.

## Conclusions

Here we describe a sustained cachexia phenotype in adult C57BL/6 mice (age 10-12 weeks) following per oral *Toxoplasma* infection. *Toxoplasma* cachexia is characterized by a loss of 20% in body mass, including fat and muscle, transient anorexia and an elevation in the hallmark cachexia cytokines IL-1, TNF and IL-6. To our knowledge, *Toxoplasma* infection is the first model to study sustained cachexia in mice that meets the modern, standard definition of cachexia put forth in 2008 (2). *Toxoplasma* infection is well established to result in acute regional ileitis, however, a detailed analysis of how long intestinal inflammation is sustained or whether acute changes in commensal microbial communities are long lived has not been asked until now. We determined that the major region of the small intestine supporting parasite replication throughout acute infection is the distal jejunum. Although we observed no evidence of sustained intestinal inflammation in chronic infection, the changes in fecal commensal communities observed in acute infection became more polarized in chronic infection. Importantly, the commensal communities of co-housed infected and uninfected mice both shifted by 5 weeks post inoculation. However, uninfected animals showed no signs of disease, suggesting that altered commensal microbiota alone is not sufficient to explain the sustained cachexia phenotype in infected animals.

Whether the cachexia program is beneficial to the host or the parasite remains to be determined. Anorexia and depletion of fat stores are classic signatures of infection that play an important role in restricting systemic bacterial pathogen replication but can trigger a host-detrimental response during viral infection(27,28). In tissue culture, the *Toxoplasma* vacuole accumulates host lipids and parasite growth can be inhibited by blocking host lipases, suggesting that the lipolysis mobilized early in infection could benefit the parasite although this has not been tested in vivo(29,30). It is now well accepted that *Toxoplasma* infection triggers altered aversion behavior to feline urine(15,16). This is hypothesized to be an adaptive strategy used by the parasite to facilitate transmission to the feline definitive host. Interestingly, the reduction in muscle mass during chronic *Toxoplasma* infection has been associated with reduced strength(22). Therefore, it is plausible that promoting cachexia during chronic infection represents a second adaptive strategy used by *Toxoplasma* to facilitate the likelihood of predation by the definitive feline host. By studying the pathways that *Toxoplasma* has evolved to manipulate to promote transmission, we may identify critical immune and metabolic interactions driving the progression of chronic cachexia that can be applied to other disease settings.

## Acknowledgements

We would like to thank Peter Buckmaster and the Stanford Veterinary Student Summer Fellowship Program for student support and constructive criticism on this project. We would like thank Erica Sonnenburg for advice and reagents for 16S sequencing.

